# A Conformational Transition of Von Willebrand Factor’s D’D3 Domain Primes It For Multimerization

**DOI:** 10.1101/2021.11.29.470312

**Authors:** Sophia Gruber, Achim Löf, Adina Hausch, Res Jöhr, Tobias Obser, Gesa König, Reinhard Schneppenheim, Maria A. Brehm, Martin Benoit, Jan Lipfert

## Abstract

Von Willebrand factor (VWF) is a multimeric plasma glycoprotein that is critically involved in hemostasis. Biosynthesis of long VWF concatemers in the endoplasmic reticulum and the (trans-)Golgi is still not fully understood. We use the single-molecule force spectroscopy technique magnetic tweezers to analyze a previously hypothesized conformational change in the D’D3 domain crucial for VWF multimerization. We find that the interface formed by submodules C8-3, TIL3, and E3 wrapping around VWD3 can open and expose two previously buried cysteines that are known to be vital for multimerization. By characterizing the conformational change at varying levels of force, we are able to quantify the kinetics of the transition and the stability of the interface. We find a pronounced destabilization of the interface upon lowering the pH from 7.4 to 6.2 and 5.5. This is consistent with initiation of the conformational change that enables VWF multimerization at the D’D3 domain by a decrease in pH in the trans-Golgi network and Weibel-Palade bodies. Furthermore, we find a stabilization of the interface in the presence of coagulation factor VIII (FVIII), providing evidence for a previously hypothesized binding site in submodule C8-3. Our findings highlight the critical role of the D’D3 domain in VWF biosynthesis and function and we anticipate our methodology to be applicable to study other, similar conformational changes in VWF and beyond.

## Introduction

Von Willebrand Factor (VWF) is a large plasma glycoprotein, critically involved in primary hemostasis. Long VWF multimers travel in the blood stream in a globular conformation and undergo conformational changes upon sensing increased hydrodynamic forces, present e.g. at sites of vascular injury^1^. Through these changes, VWF exposes binding sites for blood platelets^2,3^. After binding to collagen in the injured vessel wall, force-activated VWF thus enables formation of a hemostatic plug, built by multiple platelets binding to it^4^ (Figure 1A). The peak hydrodynamic forces acting on VWF scale approximately with the square of its length^5,6^. VWF’s occurrence in form of ultra-large concatemers, reaching lengths up to 15 μm upon unfolding^6,7^, is thus vital for effective force-activation through hydrodynamic forces at sites of vascular injury.

**Figure 1.**
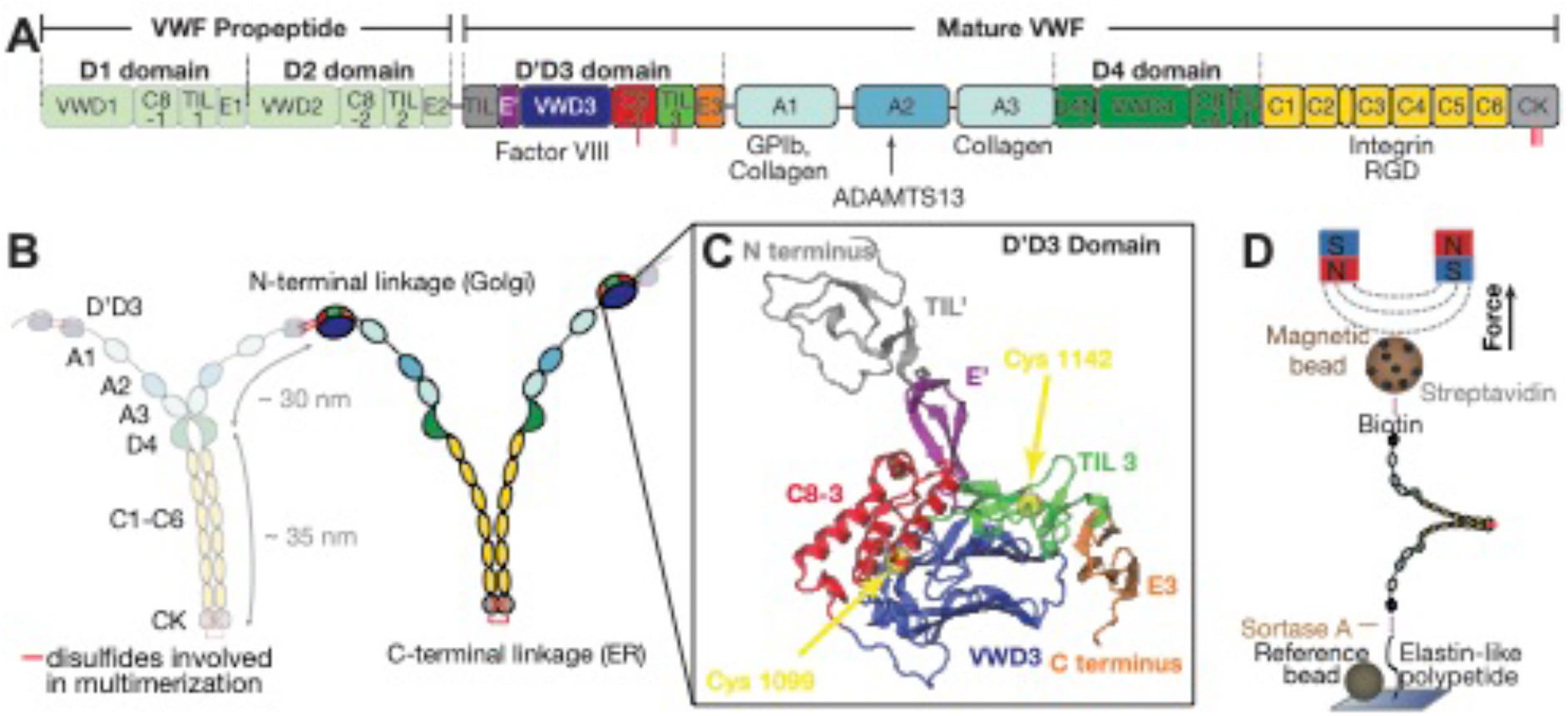
Von Willebrand Factor (VWF) domain structure and magnetic tweezers assay. **A** Domain sequence of a full VWF monomer^4^. Domains are scaled to length. The propeptide is cleaved by furin before mature VWF concatemers are secreted into the bloodstream. Binding sites of different interaction partners of VWF are indicated. **B** Mature monomers are dimerized via C-terminal linkage of the CK domains in the endoplasmic reticulum (ER) and subsequently multimerized via N-terminal linkage of two D’D3 domains in the trans-Golgi network. **C** Crystal structure of the D’D3 domain in its closed conformation (PDB accession code 6N29)^16^. The D’D3 domain comprises six submodules: TIL’ and E’ (“D’ submodules”) project out, while C8-3, TIL3, and E3 form a wedge with the larger VWD3 module (“D submodules”). Cysteines for multimerization are buried in the interface and indicated by yellow spheres. The crystal structure was rendered using VMD^42^. **D** Schematic of VWF dimer in magnetic tweezers. VWF is covalently attached to a flow cell surface via an elastin-like polypeptide (ELP) linker. Coupling to a paramagnetic bead is achieved via a stable biotin-streptavidin linkage. Reference beads are baked to the surface to account for drift. Force is applied through two permanent magnets above the flow cell.

Biosynthesis of such long concatemers is a highly complex process^8,9^: VWF is expressed as a prepropeptide, comprising a short signal peptide and the prodomains D1 and D2 in addition to the domains featured in mature VWF^10^ (Figure 1A). The signal peptide is cleaved during transport of proVWF to the endoplasmic reticulum (ER), where numerous cysteine bridges form, which shield most domains against unfolding under force^11^. In the ER, monomers dimerize via formation of three cysteine bridges between the C-terminal cystin knot (CK) domains^7,11^ (Figure 1B) and glycosylation is initiated. After dimerization, proteins are transferred to the Golgi (pH 6.2), where the stem region of VWF dimers is compacted into a “dimeric bouquet”^12^ and VWF is extensively posttranslationally modified by *N*- and *O*-glycosylation, sialylation and sulfation. In the trans-Golgi network, furin cleaves off the propeptide^4,11,13-15^, dimers assemble into a helical shape and multimerize by interdimer cysteine bonding at positions Cys1099-Cys’1099 and Cys1142-Cys’1142 in the N-terminal D’D3 domains. The multimers are stored in Weibel-Palade bodies (WPB) (secretory granules) at an even lower pH of 5.4 and secreted into the bloodstream^7,8^.

To ensure unrestricted functionality, it is of vital importance that all cysteine bridges form natively. Most disulfide bridges are formed in the ER – with the notable exception of the two cysteine bridges (Cys1099-Cys’1099 and Cys1142-Cys’1142) in the D’D3 domain mediating VWF multimerization^14^. A crystal structure of the monomeric D’D3 domain at neutral pH, characteristic of the ER, has revealed a wedge-like conformation of the D assembly^16^. In this conformation, the C8-3, TIL3, and E3 submodules make close contact with the VWD3 domain (Figure 1C), effectively burying the cysteines at positions 1099 and 1142. This conformation likely prevents premature multimerization in the ER^16^. It has been hypothesized that in the acidic pH of the (trans-)Golgi, a conformational change in the D’D3 domain exposes the cysteines to enable multimerization^16^. However, details of this necessary conformational change are currently unknown.

In addition to enabling multimerization, the D’D3 domains serve another function critical for hemostasis: By binding coagulation factor VIII (FVIII), they protect FVIII from rapid clearance^17,18^ and transport it to sites of vascular injury. Mutations within the VWF D’D3 domain that impede this high-affinity binding lead to type 2N von Willebrand disease, a condition characterized by reduced plasma levels of FVIII^19,20^. Structural and biochemical data reveal binding of the FVIII C1 domain to the D’ modules of VWF and, additionally, hint at interactions of FVIII with the VWF D3 core^16,17,21,22^.

Here, we employ magnetic tweezers (MT) to study the conformational change in the D’D3 domain necessary for multimerization as well as the interaction of the D’D3 domain with FVIII. MT are a powerful tool for single-molecule force spectroscopy, enabling multiplexed application of a large range of constant forces^23-25^. Recently, assays have been introduced employing MT for studying force-induced conformational changes in proteins^26-28^. For this purpose, single proteins are tethered between a glass surface and a magnetic bead. A magnetic field, generated by electro- or permanent magnets, exerts precisely controlled forces on the bead and thus the tethered molecule. Conformational changes in the tethered protein lead to changes in the bead position, which is monitored by video microscopy.

In this study, we investigate full-length VWF dimers under different levels of constant force (Figure 1D), to directly probe the stability of the D’D3 domain. We observe fast, reversible transitions at constant forces around 8 pN that we identify as a large-scale conformational change in the D’D3 domain (Figure 2A). Investigating the force-dependency of the transitions, we can infer the stability and dynamics of the interface. At the pH present in the Golgi and WPB, we find a significant destabilization of the interface burying the cysteines at positions 1099 and 1142 compared to neutral pH, validating the hypothesis that reduced pH plays a crucial role in VWF’s biosynthesis. Furthermore, we find a stabilization of the interface in presence of FVIII, strongly supporting the hypothesis of a binding site of FVIII within the D submodules in addition to the D’ submodules^16,21^.

**Figure 2.**
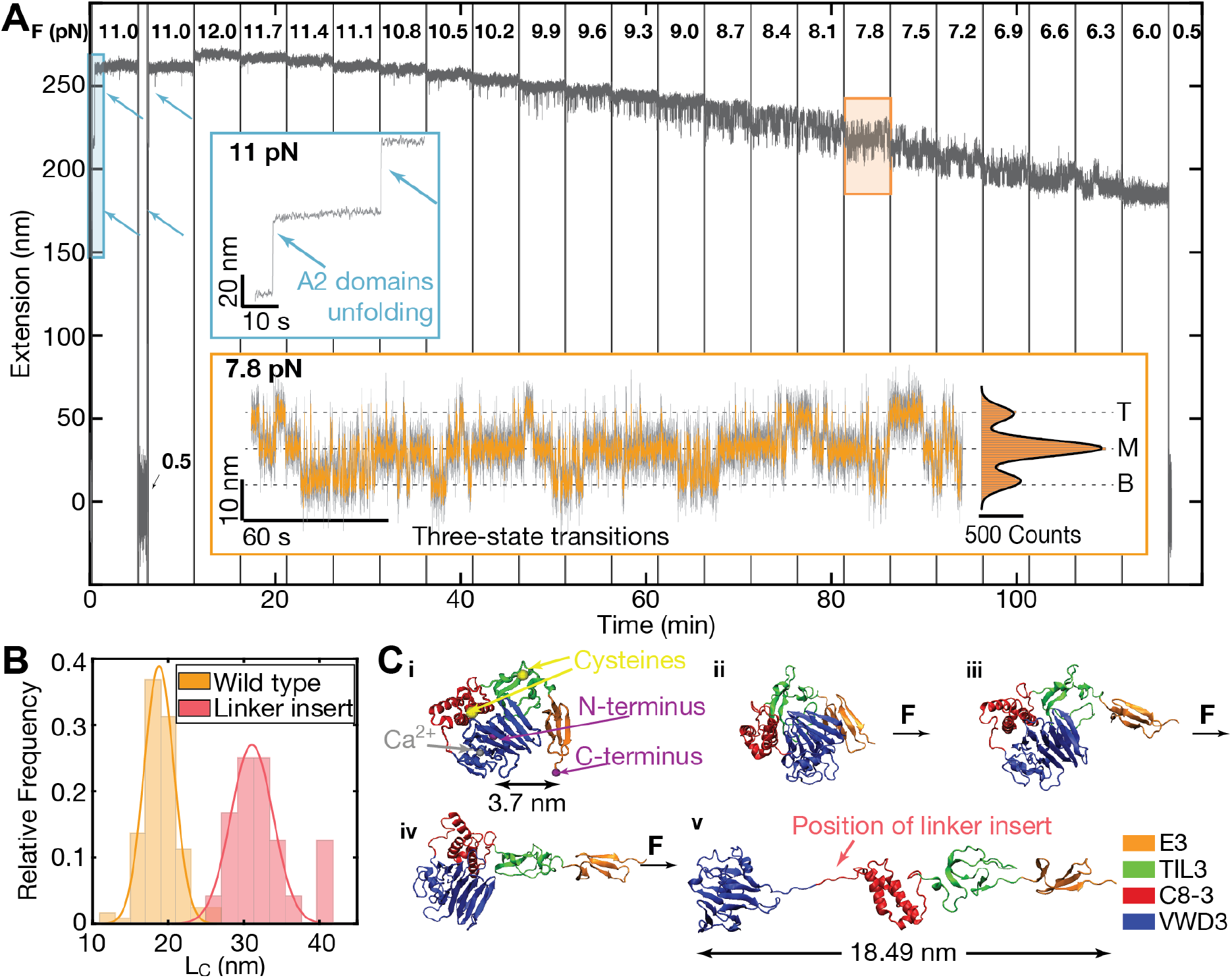
MT force spectroscopy reveals conformational change in the D’D3 domain. **A** Extension-time trace of a VWF dimer exhibiting fast, reversible transitions between three distinct states at forces around 8 pN. At the start of the measurement, two 5-min measurements at 11 pN serve to identify specific tethers by observing two ≈ 35 nm A2 unfolding events (blue inset, unfolding events indicated by arrows). Subsequently, the force is decreased from 12 pN to 6 pN in steps of 0.3 pN to systematically study the transitions between three states, separated by ≈ 7.5 nm. The population in the respective states shifts with decreasing force, with an increasing fraction of time spent in the lower extension levels at lower forces. At forces around 8 pN, transitions from the M state to the T or B state are equally likely (orange inset). In the inset, the three states (**T**op, **M**iddle, and **B**ottom) are indicated by dashed lines and are clearly visible in the extension histogram. The histogram is fit with a triple Gaussian function (black line, Table 1 Equation 10) to extract relative positions and populations of the three states. Traces recorded at 58 Hz are smoothed with an 11-frame moving average filter, and grey trace in the orange inset shows raw data. **B** Histogram of contour length transformed increment of the wild type dimer and a modified dimer with an additional 20 aa long linker insertion into the naturally occurring flexible sequence between VWD3 and C8-3 (position of the linker insert indicated in C v). Histograms were fitted with a single Gaussian (Table 1, Equation 8) using a maximum likelihood fit. Mean ± std are: L_C,wt_ = 19.0 ± 2.4 nm, L_C,linker_ = 32.4 ± 4.1 nm **C** Steered molecular dynamics (SMD) simulations validate molecular mechanism in the D’D3 domain causing transitions. **i** Crystal structure of simulated part of the D’D3 domain with VWD3 (blue), C8-3 (red), TIL3 (green), and E3 (orange)^16^. Termini are marked with purple spheres. Two cysteines involved in multimerization (Cys1099 and Cys1142) are marked with yellow spheres. Ca^2+^ is shown as a silver sphere. **ii** Initial state of SMD simulation. The pulling direction is marked with an arrow. **iii-iv** Under the influence of force, E3, TIL3, and C8-3 are “peeled” off the large VWD3 submodule. **v** Final state. Subdomain structure is kept by long-range disulfide bridges^4,16^. Total length gain in the simulation is 14.8 nm. Arrow indicates position of linker insert in red histogram in panel **B.**

## Results

To probe the stability of the wedge-like D3 interface formed by VWD3 with C8-3, TIL3, and E3, we use an assay that comprises VWF dimers, the smallest repeating subunits of long VWF concatemers (Figure 1D). We tether the dimers in MT between a flow cell surface and superparamagnetic beads using a previously described coupling strategy based on covalent surface attachment via elastin-like polypeptide (ELP) linkers^26,29^. The other terminus is attached to a superparamagnetic bead via a non-covalent, highly stable biotin-streptavidin bond^30^. In our MT setups, we apply varying levels of constant force by precisely adjusting the height of two permanent magnets located above the flow cell (Figure 1D). Among the twelve domains of mature VWF monomers, only the A2 domain is not shielded against unfolding at comparably low forces by long-range disulfide bridges. The A2 unfolding is thus one of the first responses of VWF to mechanical force and has been extensively investigated using magnetic^26^ and optical tweezers^6^ as well as atomic force microscopy-based^31^ single-molecule force spectroscopy. Here, we use the two A2 unfolding events (one from each monomer in the dimer) as a molecular fingerprint: in the beginning of each measurement, we apply a force of 11 pN and identify specifically coupled VWF dimers by observing two ≈ 35 nm steps that correspond to the A2 unfolding signature^26^ (Figure 2A, blue inset).

### Conformational Transition in the D’D3 Domain Is Revealed by MT

After selecting specific VWF tethers, we perform an inverted force ramp protocol, starting at 12 pN and decreasing the force iteratively in steps of 0.3 pN until 6 pN (Figure 2A). In the force-plateaus between 12 and 6 pN, we observe rapid, reversible transitions between a maximum of three states, named in the following **T**op, **M**iddle, and **B**ottom (Figure 2A, orange inset). The population of these states shifts with decreasing force towards the bottom state, and the midpoint force *F*_1/2_, at which transitions from the M state to the T or B state are equally likely, is found to be at around 8 pN. The three states are separated by two equidistant steps of Δz ≈ 7.6 nm at the applied forces, suggesting that the transitions stem from conformational changes that occur in each of the monomers independently. To uniquely assign the molecular origin of the observed transitions, we performed control measurements on different domain-deletion constructs, in which individual domains are deleted, but which are otherwise identical to the wild type dimer. We observe the three-state transitions independently of A2 unfolding (Figure 3A, Supplementary Figure S1A) and independently of the deletion of the D4 domain (Figure 3B, Supplementary Figure S1A) or deletion of the A1 domain (Figure 3C, Supplementary Figure S1B). Furthermore, the characteristic parameters of the transition, namely the midpoint force and the distance between the states are not altered in the domain deletion constructs (Figure 3 D, E). Therefore, we can exclude an origin within or related to the A1, A2, or D4 domains. When deleting the D’D3 domain, however, the transitions vanish (Supplementary Figure S1C), strongly suggesting that a conformational change in this domain causes the transitions.

**Figure 3.**
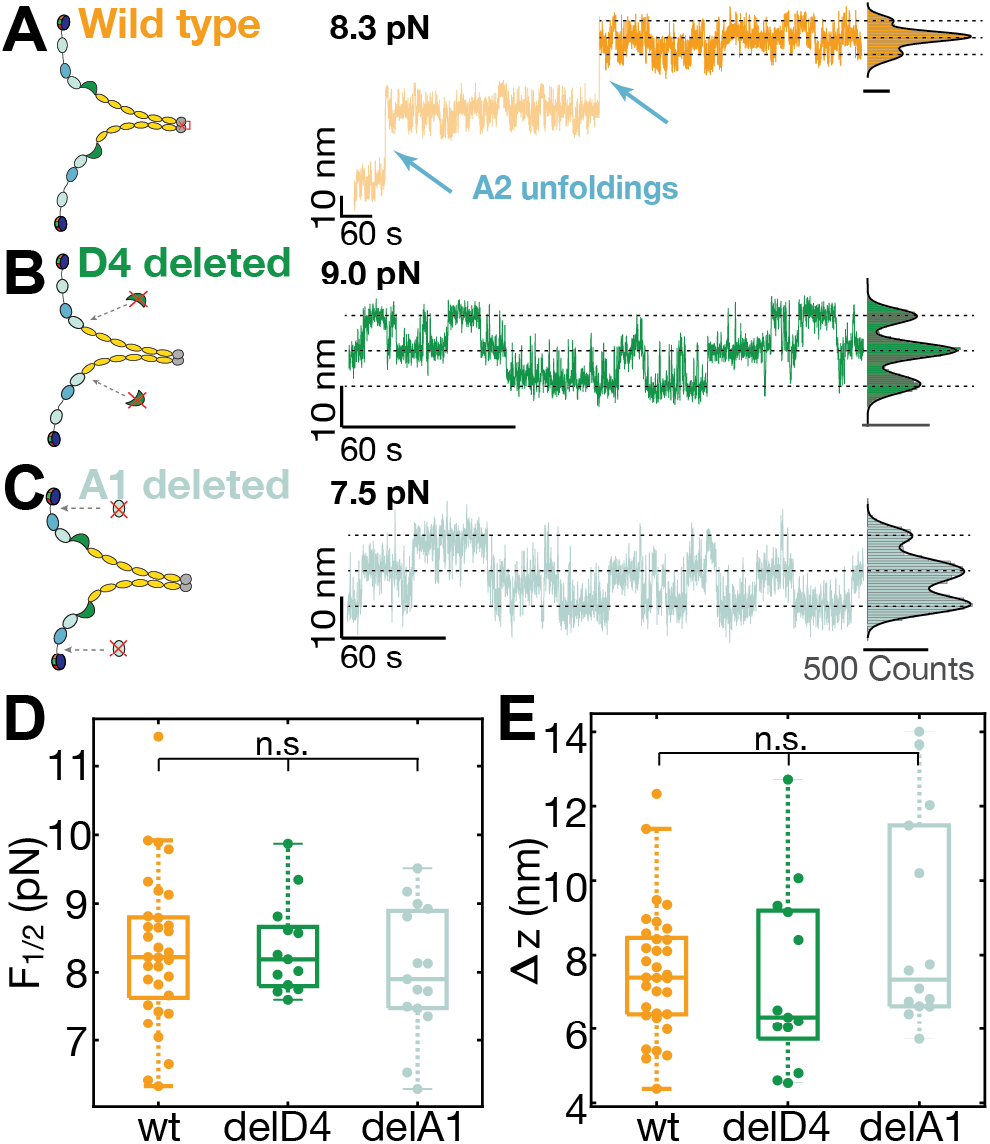
The conformational change in the D’D3 domain is unaffected by other domains. **A** Three-state transitions occur independent of the unfolding of the two A2 domains. Left: Schematic of wild type (wt) VWF dimer, right: Extension-time trace at 8.3 pN. Three-state transitions are observable before and after A2 unfolding. Extension histogram of the segment marked in dark orange reveals three distinct states that can be fitted with a three-term Gaussian (solid black line). **B** Three-state transitions occur independent of a deletion of the D4 domain. Left: Schematic of VWF dimer with deletion of both D4 domains, right: Extension-time trace at 9.0 pN. Extension histogram reveals three distinct states that can be fitted with a three-term Gaussian (solid black line). **C** Three-state transitions occur independent of a deletion of the A1 domain. Left: Schematic of VWF dimer with deletion of both A1 domains, right: Extensiontime trace at 7.5 pN. Extension histogram reveals three distinct states that can be fitted with a three-term Gaussian (solid black line). **D, E** Neither midpoint forces (D) nor Δz (E) (see main text for a discussion of the model) of the deletion constructs are significantly different from the wt. In the boxplots in D and E each data point corresponds to an individual molecule. The line in the boxes indicates the median of all data points, the box the 25^th^ and 75^th^ percentile, and the whiskers the furthest data point outside the box, but within 1.5 times the box width. Number of molecules included in D and E: wt: 33, delD4: 13, delA1: 15.

Under neutral pH and in the absence of force, the C8-3, TIL3, and E3 submodules wrap around the large VWD3, forming a wedge-like assembly that buries the two unbound cysteines^16^ Cys1099 and Cys1142 (Figure 1C). We hypothesized that by applying force, we unwrap this assembly and open the interface between VWD3, C8-3, TIL3, and E3. We verified this hypothesis by inserting 20 additional amino acids (aa) into the naturally occurring sequence between the VWD3 and C8-3 submodules (Figure 2B, C, “Linker insert”) and evaluating the distance between the states. As unfolded protein chains exhibit entropic polymer elasticity and are not completely stretched at the forces we apply, the measured distance between the states depends on the force at which the transitions occur. For the construct with the linker insert, the transitions shift to lower forces compared to the wild type construct. To ensure that the difference in force does not systematically bias the measured distance, we take into account the stretching elasticity by calculating the contour lengths from the experimentally observed distances using the worm-like chain model^26^, assuming bending persistence length *L*_p_ = 0.5 and contour length *L*_C_ = 0.4 nm per aa^26,32^. The contour length is the length of an unfolded chain of amino acids that is completely stretched and is thus independent of force. We find that inserting the 20 aa leads to an increase in contour length by 13.4 nm ± 4.8 nm (Figure 2B), in agreement, within experimental error, with the predicted 8 nm.

As an additional validation, we perform steered molecular dynamics (SMD) simulations on the D submodules (Figure 2C) by fixing the N terminus of the VWD3 submodule and pulling on the C terminus of the E3 submodule, which mimics force propagation through the D assembly in the MT assay. We find that in the simulations, the externally applied force opens the interface between VWD3 and C8-3, TIL3, and E3, while the individual subdomains initially remain folded (Figure 2C). Comparing the distance of the termini in the initial structure with their distance at the end of the simulation, we find an increase of 14.8 nm at the high forces of the SMD simulation, which is in good agreement with the experimentally determined length increase, again taking into account the protein elasticity. The results from pulling a construct with an elongated linker and the SMD simulations validate and visualize that we are opening the interface in the D3 domain in our MT assay.

### Transitions Are Inhibited in a Fraction of D3 Domains

The above findings strongly suggest that the observed transitions between three states stem from independently opening the interfaces in the two D3 domains of the tethered dimers. Interestingly, three-state transitions are not observed in all specific tethers exhibiting two A2 unfolding steps. Transitions between three states are observed in roughly 15 % of specific tethers, while in 36 %, we observe transitions only between two states, however, with otherwise identical parameters (see next section), suggesting that these correspond to conformational changes in only one of two D3 domains. The rest of the specific tethers show no transitions at all, suggesting that no conformational changes in either of the two D3 domains occur. In some cases, we initially observed three-state transitions, but in a later experiment on the same tether under the same conditions only two-state transitions. Together, these findings indicate that the conformational change can be inhibited. Considering that interface opening exposes two unbound cysteines (Figure 1C, yellow spheres), a possible explanation could be formation of non-native cysteine bridges interfering with native interface formation.

### Force-Dependent Stability of the D3 Interface

To extract thermodynamic parameters of the underlying transitions, a triple Gaussian function (Table 1, Equation 10) is fit to the extension histogram in each plateau with three-state transitions (Figure 4A, right). Thresholds are defined at the two minima between the three states (states indicated by dashed lines in Figure 4A). Based on the thresholds, the relative populations of B, M, and T states are determined at each force as the number of data points below, between, and above the thresholds divided by the total number of data points. The relative population of the three states shifts systematically with force (Figure 4B). To model the force-dependent fractions, we assume that the D3 interface can be either in a closed state or in a “peeled off’ open state. The external force biases the free energy landscape towards the open state and we assume that the free energy difference between the open and closed interface depends linearly on force as ΔG(*F*) = ΔG_0_ – *F*·Δz. Here, ΔG_0_ is the free energy difference in the absence of force and Δz is the distance between the free energy minima along the force direction. Assuming the two D3 domains in the dimer behave identically and independently, the probability of both domains being open (P_top_), one domain being open and one closed (P_middle_), and both domains being closed (P_bottom_) can be described with Equations 1-3 (Table 1). Fitting these equations to the relative population of states (Figure 4B, solid lines) yields the fit parameters F_1/2_, the midpoint force, at which it is equally likely for the domains to be open and closed, as well as Δz, the distance between the states. We find an excellent fit of the three-state model to the data (Figure 4B), which confirms the assumption of identical, independent transitions. The same analysis can be performed for molecules exhibiting two-state transitions, where only one D3 domain exhibits conformational changes (Supplementary Figure S4A). Here, a two-state model is fit to the relative state populations (Equation 4-5, Table 1), providing an independent fit of the same parameters (Supplementary Figure S4B). We find that the distributions of fit parameters obtained from analyzing molecules with two- and three-state transitions are nearly identical (Supplementary Figure S4D, E), which further supports the hypothesis that the underlying processes are indeed identical and that the intra-D3 domain transition is prevented in a fraction of D3 domains. Taking all fit parameters from two- and three-state transition molecules together (> 30 molecules; Figure 4D, E, solid lines) we find F_1/2_ = 8.3 ± 1.1 pN and Δz = 7.6 ± 1.7 nm (mean ± std), which corresponds to a mean free energy of ΔG_0_ = F_1/2_ • Δz = 9.0 ± 2.3 kcal/mol and provides a measure of the interface stability.

**Table 1:**
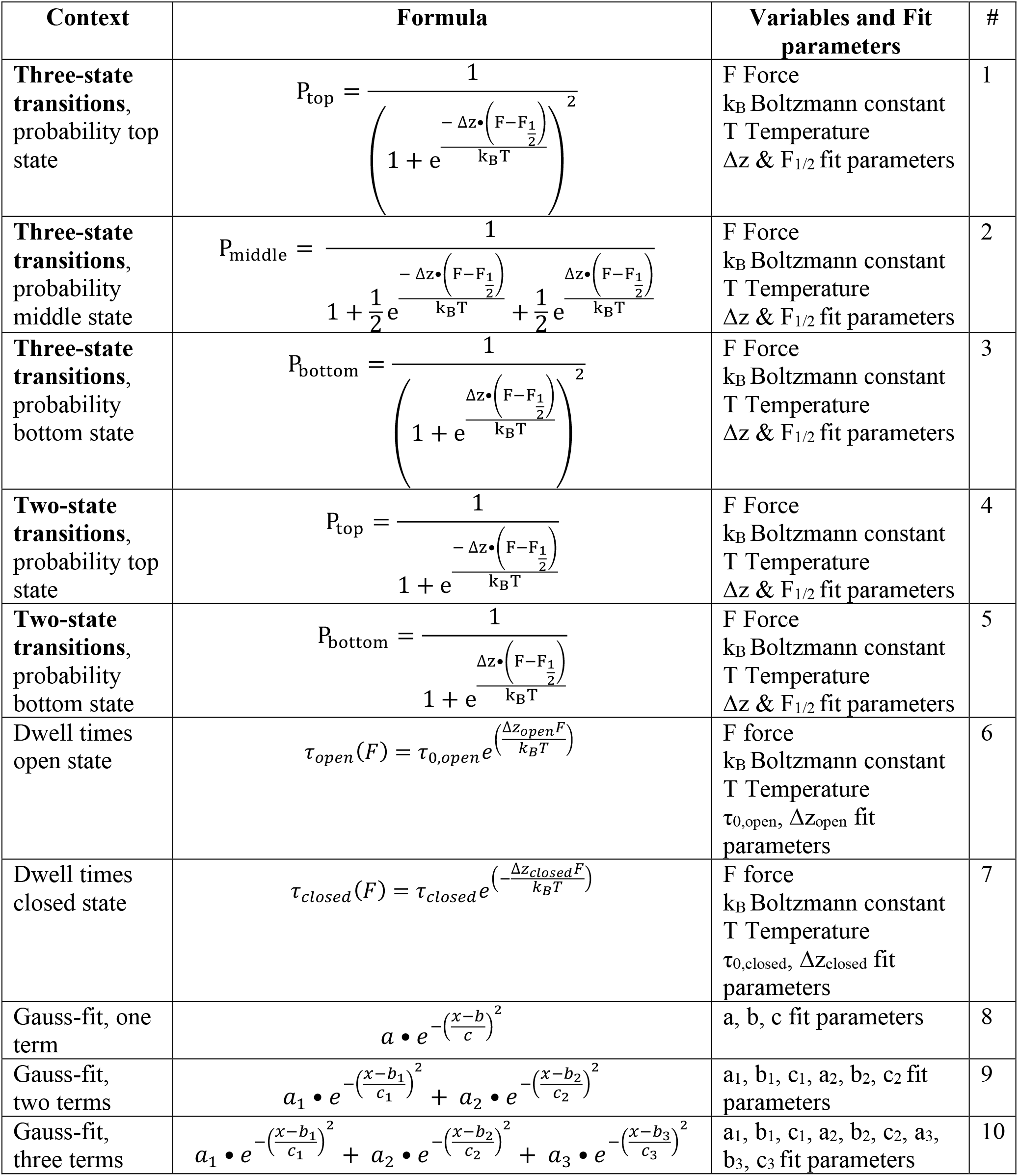
Equations and fit functions. Equations describing the equilibrium and the kinetics of the transitions in the D3 domains assuming independent processes.

| Context | Formula | Variables and Fit parameters | # |
| --- | --- | --- | --- |
| **Three-state transitions,**<br>probability top state | $P_{\text{top}} = \frac{1}{\left(1 + e^{\frac{-\Delta z \cdot \left(F - F_{1/2}\right)}{k_B T}}\right)^2}$ | F Force<br>k<sub>B</sub> Boltzmann constant<br>T Temperature<br>Δz & F<sub>1/2</sub> fit parameters | 1 |
| **Three-state transitions,**<br>probability middle state | $P_{\text{middle}} = \frac{1}{1 + \frac{1}{2} e^{\frac{-\Delta z \cdot \left(F - F_{1/2}\right)}{k_B T}} + \frac{1}{2} e^{\frac{\Delta z \cdot \left(F - F_{1/2}\right)}{k_B T}}}$ | F Force<br>k<sub>B</sub> Boltzmann constant<br>T Temperature<br>Δz & F<sub>1/2</sub> fit parameters | 2 |
| **Three-state transitions,**<br>probability bottom state | $P_{\text{bottom}} = \frac{1}{\left(1 + e^{\frac{\Delta z \cdot \left(F - F_{1/2}\right)}{k_B T}}\right)^2}$ | F Force<br>k<sub>B</sub> Boltzmann constant<br>T Temperature<br>Δz & F<sub>1/2</sub> fit parameters | 3 |
| **Two-state transitions,**<br>probability top state | $P_{\text{top}} = \frac{1}{1 + e^{\frac{-\Delta z \cdot \left(F - F_{1/2}\right)}{k_B T}}}$ | F Force<br>k<sub>B</sub> Boltzmann constant<br>T Temperature<br>Δz & F<sub>1/2</sub> fit parameters | 4 |
| **Two-state transitions,**<br>probability bottom state | $P_{\text{bottom}} = \frac{1}{1 + e^{\frac{\Delta z \cdot \left(F - F_{1/2}\right)}{k_B T}}}$ | F Force<br>k<sub>B</sub> Boltzmann constant<br>T Temperature<br>Δz & F<sub>1/2</sub> fit parameters | 5 |
| Dwell times<br>open state | $\tau_{\text{open}}(F) = \tau_{0,\text{open}} e^{\left(\frac{\Delta z_{\text{open}} F}{k_B T}\right)}$ | F force<br>k<sub>B</sub> Boltzmann constant<br>T Temperature<br>τ<sub>0,open</sub>, Δz<sub>open</sub> fit parameters | 6 |
| Dwell times<br>closed state | $\tau_{\text{closed}}(F) = \tau_{\text{closed}} e^{\left(\frac{-\Delta z_{\text{closed}} F}{k_B T}\right)}$ | F force<br>k<sub>B</sub> Boltzmann constant<br>T Temperature<br>τ<sub>0,closed</sub>, Δz<sub>closed</sub> fit parameters | 7 |
| Gauss-fit, one term | $a \cdot e^{-\left(\frac{x-b}{c}\right)^2}$ | a, b, c fit parameters | 8 |
| Gauss-fit, two terms | $a_1 \cdot e^{-\left(\frac{x-b_1}{c_1}\right)^2} + a_2 \cdot e^{-\left(\frac{x-b_2}{c_2}\right)^2}$ | a<sub>1</sub>, b<sub>1</sub>, c<sub>1</sub>, a<sub>2</sub>, b<sub>2</sub>, c<sub>2</sub> fit parameters | 9 |
| Gauss-fit, three terms | $a_1 \cdot e^{-\left(\frac{x-b_1}{c_1}\right)^2} + a_2 \cdot e^{-\left(\frac{x-b_2}{c_2}\right)^2} + a_3 \cdot e^{-\left(\frac{x-b_3}{c_3}\right)^2}$ | a<sub>1</sub>, b<sub>1</sub>, c<sub>1</sub>, a<sub>2</sub>, b<sub>2</sub>, c<sub>2</sub>, a<sub>3</sub>, b<sub>3</sub>, c<sub>3</sub> fit parameters | 10 |

**Figure 4.**
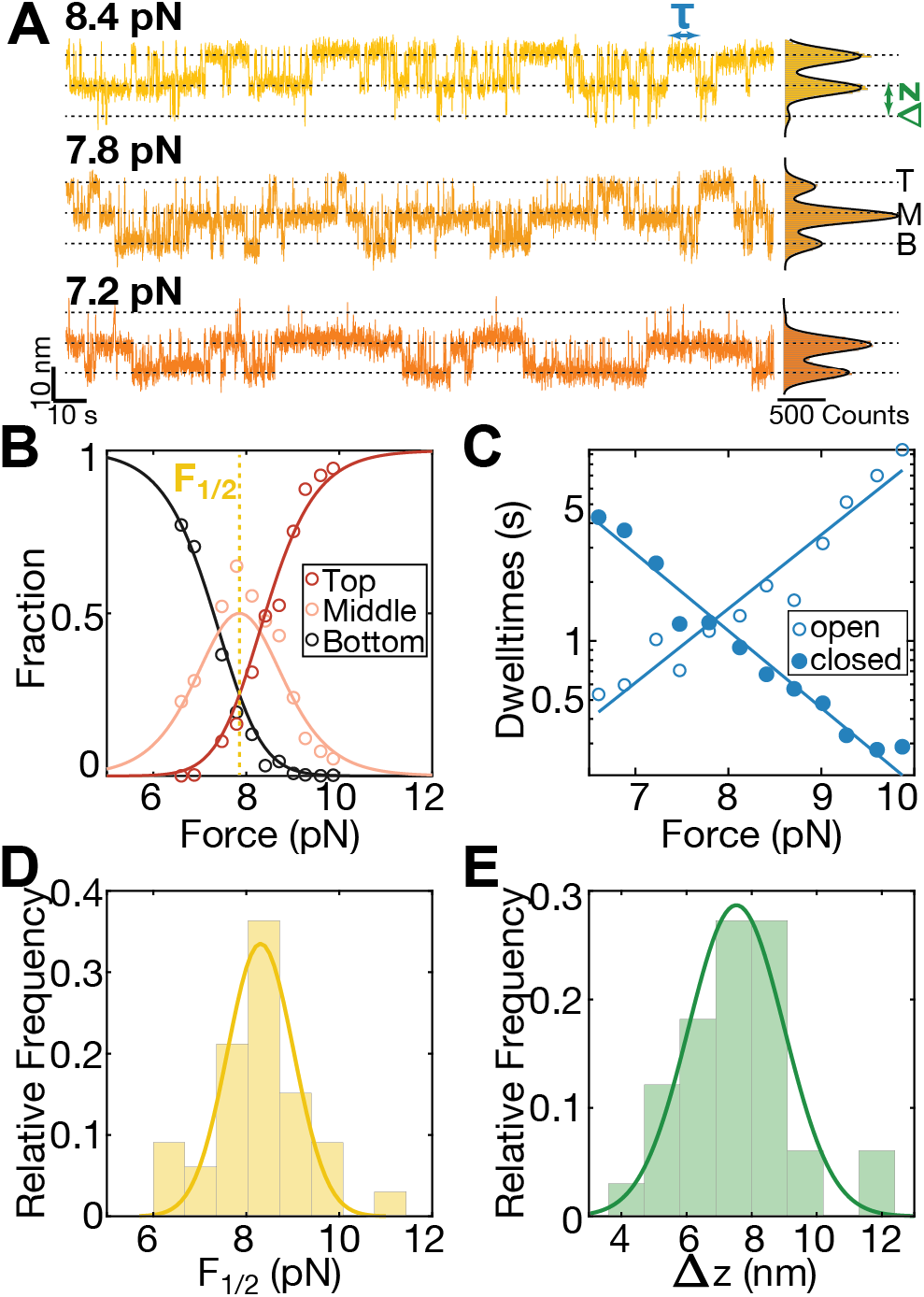
Stability and dynamics of the D3 interface probed by MT force spectroscopy. **A** Extension-time traces of VWF dimers in MT. Three-state transitions shift with decreasing force towards the bottom state. The three states (**T**op, **M**iddle, and **B**ottom) are indicated by dashed lines. Histograms of the extensions are shown on the right. Black lines show triple Gaussian fits. **B** Analysis of the relative population of the different states as a function of force. Circles represent experimental data with each circle corresponding to a 5-min force plateau. A three-state model assuming two independent transitions fits the experimental data well (solid lines, Table 1, Equations 1-3). Fit parameters are the mid-force F_1/2_ (here F_1/2_ = 7.85 pN), at which domains are equally likely open or closed, and the distance between the states Δz (here Δz = 6.5 nm). **C** Pseudo dwell time distributions. Pseudo dwell times for open and closed domains are calculated from dwell times in the top, middle, and bottom state. For each plateau, pseudo dwell times are determined and fit with an exponential to determine the mean dwell times for each force. Mean dwell times in the open (open circles) and the closed state (filled circles) depend exponentially on the applied force (solid lines are exponential fits, Table 1, Equations 6-7). **D** Histogram of mean midpoint forces determined from fitting the two or three-state model (panel B, Supplementary Figure 4B). The solid line is a Gaussian fit with mean ± std: 8.3 ± 1.1 pN. **E** Histogram of Δz determined from fitting the two or three-state model (panel B, Supplementary Figure 4B). The solid line is a Gaussian fit with mean ± std: 7.6 ± 1.7 nm. Histograms in **D** and **E** show distributions of 33 molecules.

### Kinetics of the Conformational Changes in the D3 Interface

In addition to providing insights into the force-dependent equilibrium, our MT extension time traces can reveal kinetic information from the transitions at different forces. Using the same thresholds as for state-population analysis, we identify dwell times (Figure 4A, τ; Supplementary Figure S3A, B) as times that are spent in one state before crossing the threshold. The dwell times in the three-state transitions, however, reflect the kinetics of two equal processes happening independently at the same time. To access the dwell times of individual domains, τ_open_ and τ_closed_, dwell times in the middle plateau are divided by two and associated with the dwell times in the bottom state and the top state, respectively. This procedure to obtain so-called *pseudo-dwell times* for individual domains takes into account the number of domains opened and closed in each state and weighs the measured dwell times accordingly^33,34^. We find the pseudo-dwell times for each plateau to be exponentially distributed (Supplementary Figure S3C, D). The force-dependent mean pseudo dwell times are well-described by exponential, Arrhenius-like relationships^35^ (Table 1, Equations 6 and 7; Figure 4 C, solid lines). Fitting parameters τ_0, open_ and τ_0, closed_ are the lifetimes of the open and closed conformation at zero force and Δz_open_ and Δz_closed_ are the distances to the transition state along the pulling direction. The sum of Δz_open_ and Δz_closed_ (3.1 nm + 4.7 nm = 7.8 nm) is found to be in excellent agreement with Δz obtained from fitting relative state populations, which provides a consistency check between equilibrium and kinetic analysis and suggests that there is a single dominant energy barrier along the reaction pathway. The extrapolated lifetimes at zero force of the closed conformation τ_0, closed_ are in the range of hours. In comparison, the lifetimes of the open states in the absence of load τ_0, open_ are much shorter, only on the order of milliseconds. The extrapolated lifetimes at zero force provide another route for calculating the mean free energy: ΔG_0,τ_ = k_B_T • log(τ_0,open_/τ_0,closed_) = 9.3 ± 1.7 kcal/mol, in excellent agreement with the ΔG_0_ value computed from the force-dependent populations in the previous section (Figure 5C). The good agreement between the free energy differences obtained from equilibrium and lifetime analysis provides another consistency check between the thermodynamic and kinetic analyses.

**Figure 5.**
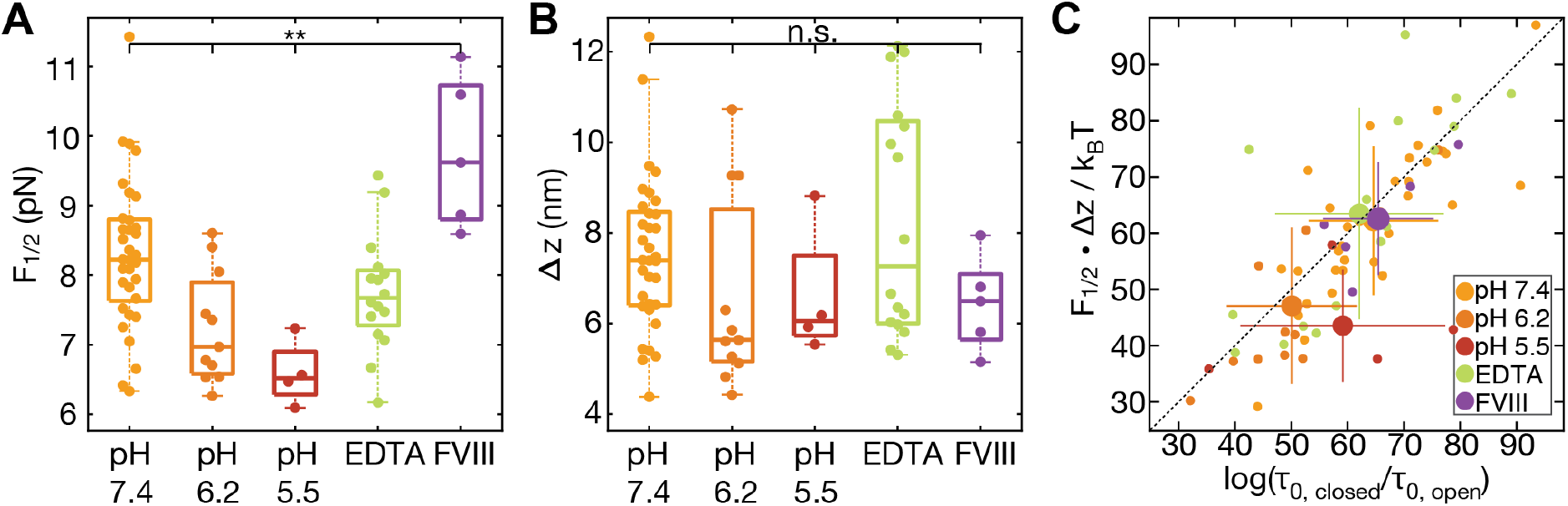
The stability of the D3 interface is modulated by pH and FVIII binding. **A** Midpoint force F1/2 for different buffer conditions (pH 7.4 [yellow]; pH 6.2 [orange]; pH 5.5 [red]; pH 7.4 + EDTA [green]; pH 7.4 + FVIII [purple]). Buffer compositions are listed in Supplementary Table 1. The D3 interface is destabilized by decreasing pH, not affected by addition of EDTA (to remove the Ca^2+^ from the binding loop in VWD3), and stabilized by addition of FVIII. The difference between F1/2 at pH 7.4 and pH 6.2 is highly significant at p < 0.00412 (two-tailed t test for two independent means). The difference between F1/2 at pH 7.4 and pH 7.4 + FVIII is highly significant at p < 0.00781 (two-tailed t test for two independent means). **B** There are no significant differences in Δz values under all conditions. In the boxplots in A and B each data point corresponds to an individual molecule. The line in the boxes indicates the median of all data points, the box the 25th and 75th percentile, and the whiskers the furthest datapoint outside the box, but within 1.5 times the box width. **C** Free energy differences between the open and closed state of the D3 interface. The free energy differences were obtained from the equilibrium data as F1/2•Δx and from the kinetics as k_B_T•log(τ_0, closed_/τ_0,open_). The data fall along the 45° line (dashed), indicating that the two estimates give consistent values. Comparison of the different conditions reveals a lower free energy difference for decreased pH of 6.2 and 5.5, indicating a destabilization of the domain at low pH. Number of molecules for the five conditions: pH 7.4: 33; pH 6.2: 11; pH 5.5: 4; pH 7.4 + EDTA: 16; pH 7.4 + Factor 8: 5.

### Lowering pH Destabilizes the D3 Interface

We analyzed the equilibrium and kinetics of the conformational change under different physiologically relevant conditions (Figure 5). Specifically, we compared pH 7.4, characteristic of blood, with the lower pH conditions of pH 6.2 and 5.5, representative of the conditions in the trans-Golgi network and WPB, respectively. We find that the extension change Δz is insensitive to pH (Figure 5B), suggesting that the overall fold of the D3 domain and geometry of the transition is not affected in this pH range. However, we find a significant destabilization of the interface (p < 0.00412; two-tailed t test for two independent means, performed on F_1/2_), reflected both in a lower midpoint force (Figure 5A, yellow [pH 7.4]: 8.3 ± 1.1 pN, orange [pH 6.2]: 7.2 ± 0.8 pN, and red [pH 5.5]: 6.6 ± 0.5 pN), and a corresponding decrease in ΔG_0_ (Figure 5C, pH 7.4: 9.0 ± 2.3 kcal/mol, pH 6.2: 6.8 ± 2.3 kcal/mol, pH 5.5: 6.3 ± 1.5 kcal/mol). Furthermore, the D3 interface becomes more dynamic and in particular the extrapolated lifetime in the open conformation is increased by decreased pH from 0.004 s at pH 7.4 to ~0.3 s at pH 6.2 and pH 5.5, suggesting an approximately 10-fold higher exposure of the cysteines buried by the D3 interface at acidic pH. Previously, it was reported that for successful VWF multimerization low pH and Ca^2+^ are required^7^. To investigate the role of Ca^2+^, we performed measurements in the presence of 10 mM EDTA, to chelate divalent ions. In contrast to decreased pH, we found that EDTA does not significantly affect the stability of the interface (Figure 5A-C). The fact that the addition of EDTA does not alter the stability of the interface could either indicate that Ca^2+^ ions have only limited influence on the interface or that structural ions, e.g. the ion positioned in a Ca^2+^ binding loop in VWD3 (Figure 2C, i), are so stably bound that they are not efficiently removed by EDTA.

### FVIII Binding Stabilizes the D3 Interface

We next used our MT assay to probe the transitions in the D3 domain in the presence of FVIII. We find a highly significant stabilization (p < 0.00781; two-tailed t test for two independent means, performed on F_1/2_) of the D3 interface in the presence of ≈ 640 nM FVIII. FVIII has been reported to predominantly bind to the D’ submodules^21,22^. Based on point mutations in the C8-3 domain, which lead to 2N von Willebrand disease phenotypes with decreased ability to bind FVIII, it was however hypothesized that FVIII could also have a less prominent interaction site in the C8-3 module^16^. Our results strongly support a binding site in the D3 domain, as FVIII directly impacts the conformational change of the D3 interface.

### Absence of D3 Interface Transitions in Multimerized D’D3 Domains

There is currently no crystal structure available for a dimerized D’D3 domain. Therefore, the native force propagation through the D’D3 domain in multimerized VWF is not known. To assess whether the conformational change in the D3 domain would only play a role in biosynthesis or might also occur in VWF multimers and potentially be relevant for VWF force-activation, we investigated an “inverted” VWF dimer that is dimerized via its D’D3 domains (Supplementary Figure S5). Here, we again observed A2 unfolding events, but no additional transitions, suggesting that the transition in the D3 interface only occurs prior to multimerization at the interface.

## Discussion

VWF multimerization is a crucial process for successful hemostasis. It is known that the free cysteines Cys1099 and Cys1142, located in the N-terminal D’D3 domains, are crucially involved in multimerization in the trans-Golgi. However, the crystal structure of a monomeric D’D3 domain at neutral pH shows these two cysteines buried in a wedge-like structure formed by the VWD3 – C8-3, TIL3, and E3 interface. The details of how the cysteines are exposed to enable multimerization were previously unknown. Recently, Springer and coworkers hypothesized that Cys1099 attacks the cysteine bond between Cys1097 and Cys1091, forming a new bond with Cys1091 and releasing Cys1097 for disulfide bond formation with Cys1097’ in a second dimer^36^. However, no matter whether Cys1099 or Cys1097 forms an inter-dimer disulfide bond alongside Cys1142, there has to be a prior conformational change to expose both cysteines buried in the wedge. It has been hypothesized that this conformational change is induced by the acidic pH in the Golgi apparatus^16,36^. Here, we used MT to examine a conformational transition in the D3 domain, peeling submodules C8-3, TIL3, and E3 off the larger VWD3 submodule and thus exposing the unbound cysteines. Studying the D3 interaction under various levels of constant force, we obtained force-dependent populations and rates that we extrapolated to zero load to characterize the stability of the interface. Both the extrapolated lifetimes and the estimated free energies suggest that the D3 interface is very stable at neutral pH, with a fraction of only about 1 in 10^7^ molecules being in the open conformation at any given time in the absence of force, thus effectively shielding the free cysteines buried by the interface. At lower pH, characteristic of (trans-)Golgi and WPB, the D3 interface is significantly destabilized and becomes more dynamic. The pronounced pH dependency can be rationalized on the molecular level by the large number of histidine-residues in the interface that can be protonated at acidic pH and then likely destabilize the interaction between the four submodules^16^. Biologically, the regulation of the conformational change by pH is of great importance for VWF’s biosynthesis to enable exposure of the buried cysteines involved in multimerization only under the acidic pH in the trans-Golgi network. The stability of the interface revealed by our assay suggests that VWF dimers are protected from forming premature cysteine bridges involving Cys1099 and Cys1142 in the ER, which would have the potential to disturb organized compaction and multimerization in the trans-Golgi network. Finally, we characterized the D3 interface opening in the presence of FVIII. We found a statistically significant stabilization of the interface, indicating that FVIII not only binds to the D’ modules, but also to the D modules of the D’D3 domain, as had been suggested by structural and biochemical information. Our results highlight how complex interactions regulate the biosynthesis and function of VWF and demonstrate how MT force spectroscopy can probe biologically relevant conformational changes under a broad range of conditions.

## Materials and Methods

### VWF Constructs

Dimeric VWF constructs were designed as hetero-bifunctional dimers, consisting of two different types of monomers possessing different N-terminal peptide tags. One monomer possessed a ybbR-tag, allowing for covalent conjugation of CoA-biotin. The second monomer was equipped with a strep-tag II for high-affinity purification, followed by a tobacco etch virus (TEV) protease cleavage site^37^ and the N-terminal sortase motif^38^ GG. The TEV site served two purposes: First, to remove the strep-tag after purification, to avoid interaction with streptavidin on the magnetic beads, and second, to expose the sortase motif GG, which must be located terminally for the sortase-mediated ligation to the ELP linker. In addition to full-length dimers, comprising all domains present in mature VWF, also several constructs with deletions of certain domains were investigated as controls: delD4, with a deletion of the full D4 assembly (D4N-TIL4, aa 1873-2255), delD’D3, with a deletion of the full D’D3 assembly (TIL’-E3, aa 764-1273), and delA1, with a deletion of the A1 domain (aa 1272-1462). Additionally, an “inverted” construct was expressed, which dimerized N-terminally. To produce such N-terminally, but not C-terminally linked dimers, monomers with mutation p.Cys2771Arg in the CK domain to impair C-terminal dimerization^39,40^ and with C-terminal tags for site-specific protein attachment were expressed in the presence of the VWF propeptide (VWFpp) domains D1 and D2.

Hetero-bifunctional dimers were obtained by co-transfection of HEK-293 cells with two different plasmids so that the two different types of monomers were co-expressed. Multimerization was obstructed by deleting the VWF pro-peptide sequence (domains D1 and D2, aa 26-763). N-terminal tags were inserted after the required N-terminal signal peptide (aa 1-25). Plasmid construction, transfection of HEK-293 cells and protein expression were performed as described in detail in Müller *et al*.^31^. In brief, 2•10^6^ HEK-293 cells (DSMZ, Germany) were transfected in Dulbecco’s modified Eagle’s medium (Life Technologies) containing 10 % fetal bovine serum (Life Technologies), 2 μg of each of the two plasmids, and 15 μl Lipofectamine 2000 (Life Technologies).

24 h after transfection, cells were transferred into selection medium containing 500 μg/ml G418 (Invivogen) and 250 μg/ml Hygromycin B (Invivogen). After 2–3 weeks, the polyclonal cell culture was seeded for expression. After 72 h of cell growth, the medium was exchanged against OPTIPRO serum-free medium (Life Technologies) for serum-free collection of secreted recombinant VWF. The culture supernatant was collected after 72 h and concentrated using Amicon Ultra-15 MWCO 100 kDa (Merck Millipore).

All dimeric constructs were purified via a HiTrap StrepTrap affinity chromatography column (GE Healthcare) using the AEKTA Explorer system (GE Healthcare). As running buffer, 20mM Hepes, 150mM NaCl, 1mM MgCl_2_, 1mM CaCl_2_, pH 7.4, was used. Elution buffer additionally contained 2.5mM d-desthiobiotin. Eluted VWF constructs were buffer exchanged (to the running buffer) and concentrated by centrifuge filtration using Amicon UltraMWCO 100 kDa (Merck Millipore). After 5 washing steps with 500 μl of running buffer, the volume was decreased to ≈ 100 μl before inverting the filter and harvesting the protein.

### Steered Molecular Dynamics Simulations

The steered molecular dynamics (SMD) simulations were performed using the crystal structure of the monomeric von Willebrand Factor D’D3 assembly by Dong *et al*. (PDB-ID 6n29)^41^. Further structure preparation as well as the MD simulations were performed using VMD with the QwikMD plugin^42,43^. Standard parameters defined by QwikMD have been used for the simulations. The MD simulations were conducted using the NAMD molecular dynamics package^44^ together with the CHARMM36 force field^44,45^.

The disulfide bonds that preserve the structure of the individual subdomains were treated using the classical MD forcefields parameters. As a first pre-processing step, the original crystal structure was solvated in a box containing TIP3 water molecules and a NaCl concentration of 0.15 M. Before the pulling experiment, the protein was relaxed by molecular dynamics simulation for 100 ns. For the steered molecular dynamics experiment, the water box was enlarged in the pulling direction and the molecule was pulled at the C-terminal glycine while the N-terminal aspartate was anchored SMD simulations were run with an applied pulling speed of 1 Å/ns for 150 ns.

### MT Instrument

MT experiments were performed on a previously described custom setup^26,46^. The setup employs a pair of permanent magnets (5×5×5 mm^3^ each; W-05-N50-G, Supermagnete, Switzerland) in vertical configuration^47^. The distance between magnets and flow cell (and, thus, the force) is controlled by a DC-motor (M-126.PD2; PI Physikinstrumente, Germany). An LED (69647, Lumitronix LED Technik GmbH, Germany) is used for illumination. A 40x oil immersion objective (UPLFLN 40x, Olympus, Japan) and a CMOS sensor camera with 4096×3072 pixels (12M Falcon2, Teledyne Dalsa, Canada) allow to image a large field of view of approximately 440 × 330 μm^2^ at a frame rate of 58 Hz. Images are transferred to a frame grabber (PCIe 1433; National Instruments, Austin, TX) and analyzed with a LabView-based open-source tracking software^48^. The bead tracking accuracy of the setup is ≈ 0.6 nm in (x, y) and ≈ 1.5 nm in z direction. For creating the look-up table required for tracking the bead positions in z, the objective is mounted on a piezo stage (Pifoc P-726.1CD, PI Physikinstrumente). Force calibration was conducted as described by te Velthuis *et al*.^49^ based on the transverse fluctuations of long DNA tethers. Importantly, for the small extension changes on the length scales of our protein tethers, the force stays essentially constant^26^, with the relative change in force due to tether stretching or protein unfolding being < 10^-4^. Force deviations due to magnetic field inhomogeneities across the full range of the field of view are < 3%. The largest source of force uncertainty is the bead-to-bead variation, which is on the order of ≤ 10% for the beads used in this study^26,47,50,51^.

### Single Molecule MT Measurements

Preparation of flow cells was performed as described^26^. In brief, aminosilanized glass slides were functionalized with elastin-like polypeptide (ELP) linkers^52^, possessing a single cysteine at their N terminus as well as a C-terminal Sortase motif, via a small-molecule crosslinker with a thiol-reactive maleimide group [sulfosuccinimidyl 4-(N-maleimidomethyl)cyclohexane-1-carboxylate; Sulfo-SMCC, Thermo Fisher Scientific]. Flow cells were then assembled from an ELP-functionalized slide as bottom and a nonfunctionalized glass slide with two small holes for inlet and outlet as top, with a layer of cutout parafilm (Pechiney Plastic Packaging Inc., Chicago, IL) in between to form a channel. Flow cells were incubated with 1 % casein solution (Sigma-Aldrich) for 1 h and flushed with 1 ml (approximately 20 flowcell volumes) of buffer pH 7.4, near physiological (Supplementary Table 1).

CoA-biotin (New England Biolabs) was coupled to the ybbR-tag of the VWF-dimer constructs in a bulk reaction in the presence of 5 μM sfp phosphopantetheinyl transferase^53^ and 10 mM MgCl_2_ at 37 °C for 60 min. Afterwards, VWF dimers were diluted to a final concentration of approximately 20 nM in pH 7.4, near physiological buffer (Supplementary Table 1), and incubated in the flow cell in the presence of 2 μM evolved pentamutant Sortase A^54,55^ for 30 min. Subsequently, the flow cell was flushed with 1 ml of measurement buffer (pH 7.4, near physiological buffer with 0.1 % Tween-20 (Supplementary Table 1)). Finally, beads functionalized with streptavidin (Dynabeads M-270, Invitrogen) were incubated in the flow cell for 60 s, and unbound beads were flushed out with 1 ml of measurement buffer.

At the beginning of each measurement, the tethered beads were subjected to two 5-min intervals of a constant force of 11 pN to allow for identification of specific, single-tethered beads by the characteristic unfolding of the two A2 domains^26^ (Fig. 2A, green inset). Only beads that showed two A2 unfoldings were analyzed further. After 30 s at a low resting force of 0.5 pN, beads were subjected to a forceramp starting at 12 pN and going down to 6 pN in steps of 0.3 pN, with each plateau of constant force lasting for 5 minutes. All measurements were performed at room temperature ≈ 22 °C). For testing the transitions under different buffer conditions, buffer was exchanged and the same measurement protocol was repeated. Buffer composition of different buffers are listed in Supplementary Table 1. To test the influence of Factor VIII, we used a haemophilia A drug – Advate 500 I.E. (Baxter AG, Wien, Austria) – containing octocog alfa, recombinant human factor VIII. After diluting the powder according to the manufacturers instructions, we filtered the recombinant F-VIII, exchanged the buffer to our measurement buffer with pH 7.4, and increased the concentration by centrifuge filtration using Amicon UltraMWCO 100 kDa (Merck Millipore): After 5 washing steps with the measurement buffer, we decreased the volume to ≈ 90 μl before harvesting F-VIII by filter inversion. This resulted in a F-VIII concentration of 0.642 μM. The ≈ 90 μl were inserted into the flowcell and incubated for 30 min before repeating the same measurement protocol described above.

### Data Analysis

Data analysis was performed with custom MATLAB scripts. Specific VWF dimer tethers were selected on the basis of the two A2 fingerprint unfolding events, characterized previously^26,31^. Extension vs. time traces were subjected to a tether-specific smoothing with a moving average filter. The number of frames used for the smoothing was determined based on the distance of the states and the Allan Deviation (AD) of the tether extension fluctuation (see Supplementary Figure 2). The distance between the states Δz was determined from fitting a double or triple Gaussian function to the histogram of the extension in the lowest constant force plateau exhibiting a population in all two or three-states, respectively and evaluating the distance between the peaks, as shown in Supplementary Figure 2B to the left (1.)). The AD was calculated from a 30 second fragment of the trace at the lowest constant force plateau exhibiting no transitions and fit with a theoretical model of an overdamped bead (Supplementary Figure 2B middle (2.))). The AD is defined as

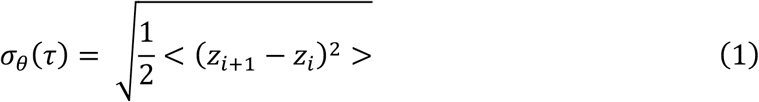

where z_i_ is the mean of the measurement interval of length τ. The angle bracket denotes the arithmetic mean over all measurement intervals. In other words, the AD is one-half of the averaged square distance between the means of neighbouring intervals^56-58^. Intuitively, it gives a measure of the spatial resolution after averaging over a time interval τ. Using the criterion that the deviation should be at least four times smaller than the evaluated distance Δz, a smoothing factor for each trace can be determined by multiplication of the averaging interval τ, where AD equals Δz/4, with the measurement frequency. This smoothing factor – typically in the range of 8 to 13 frames – is applied to the time extension-time trace before evaluation of the state-population and the dwell times in each force plateau.

## Authors Contributions

A.L., S.G., T.O., M.A.B., M.B., and J.L. designed research; S.G., and A.H. built instruments; A.L., S.G., and A.H. performed experiments; R.J. performed and analyzed simulations; T.O. and G.K. expressed proteins; A.L., and S.G. analyzed experimental data; and A.L., S.G., and J.L. wrote the paper with input from all authors.

## Acknowledgements

We thank Thomas Nicolaus and Angelika Kardinal for laboratory assistance, Wolfgang Ott for providing ELP linkers, Magnus Bauer, Rafael C. Bernadi, Philipp U. Walker, Hermann E. Gaub, Frauke Gräter, Philip J. Hogg, Camilo Aponte-Santamaría, and Fabian Kutzki for helpful discussions. This project was funded by the Deutsche Forschungsgemeinschaft (DFG, German Research Foundation) Project-ID 201269156, SFB 1032 and Project-ID 386143268, “Unraveling the Mechano-Regulation of Von Willebrand Factor”.

## Supplementary Material

### Supplementary Figures

**Supplementary Figure S1.**
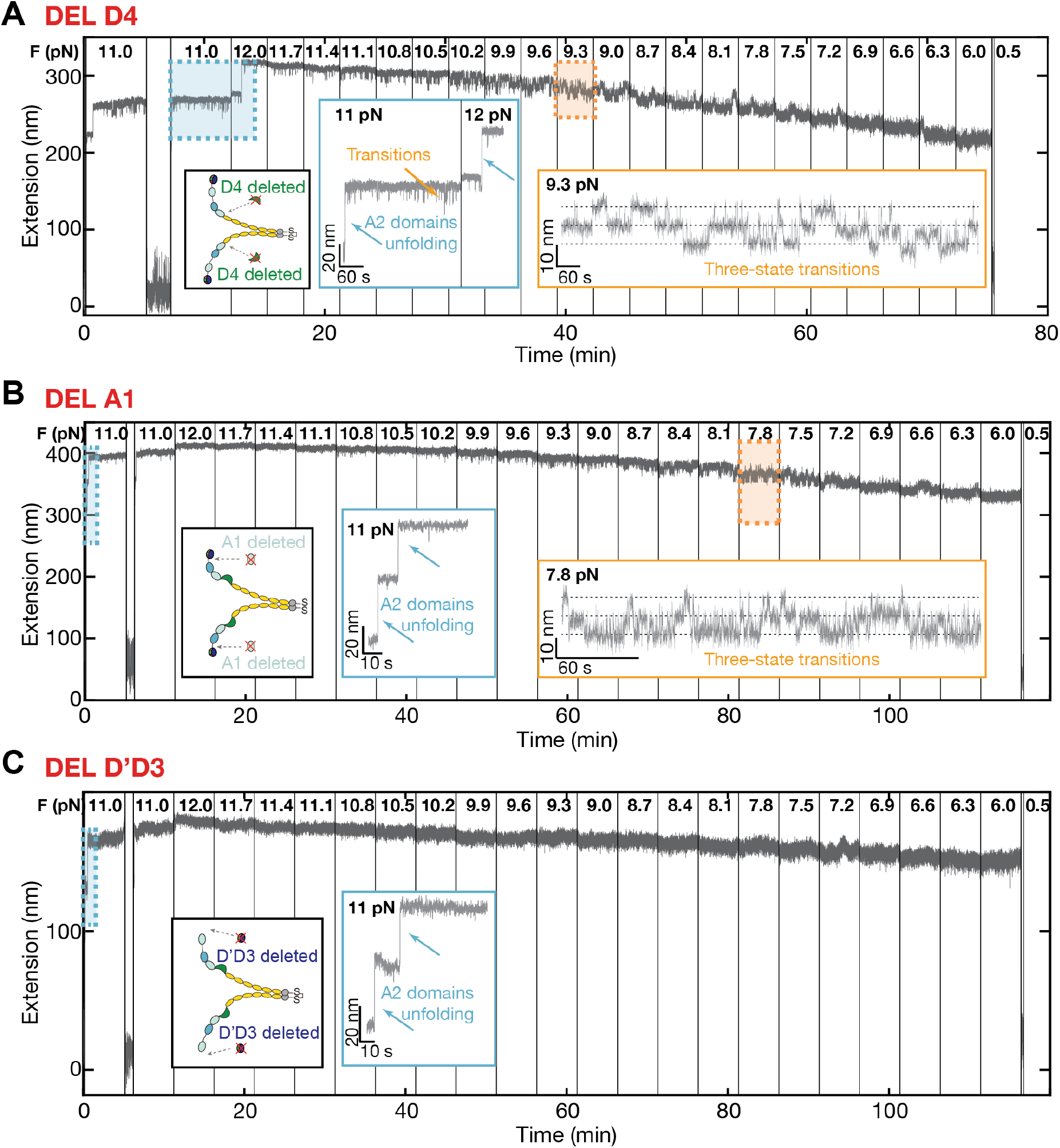
Control measurements with domain deletions to assign the transitions to the D’D3 domain. Individual domains were deleted to verify the origin of the transitions in the D’D3 domain. Apart from deleted domains, constructs possessed the same tags and were objected to the same force protocol as the wild type. **A** Heterodimer with D4 domain deletions (schematic of construct shown as inset (black frame)). Molecules with D4 domain deletions show A2 domain unfolding events (inset with blue frame) and three-state transitions (inset with orange frame), proving transitions to be independent of the D4 domain. Notably, transitions also appear in between the two A2 unfolding events, suggesting that transitions are also independent of the A2 domain. **B** Heterodimer with A1 domain deletions (schematic of construct shown as inset (black frame)). Molecules with A1 domain deletions show A2 domain unfolding events (inset with blue frame) and three-state transitions (inset with orange frame), proving transitions to be independent of the A1 domain. **C** Heterodimer with D’D3 domain deletions (schematic of construct shown as inset (black frame)). Constructs with D’D3 domain deletions show A2 domain unfolding events, but no transitions. This was checked for > 40 molecules. The absence of transitions in this deletion construct strongly suggests that transitions originate in the D’D3 domain.

**Supplementary Figure S2.**
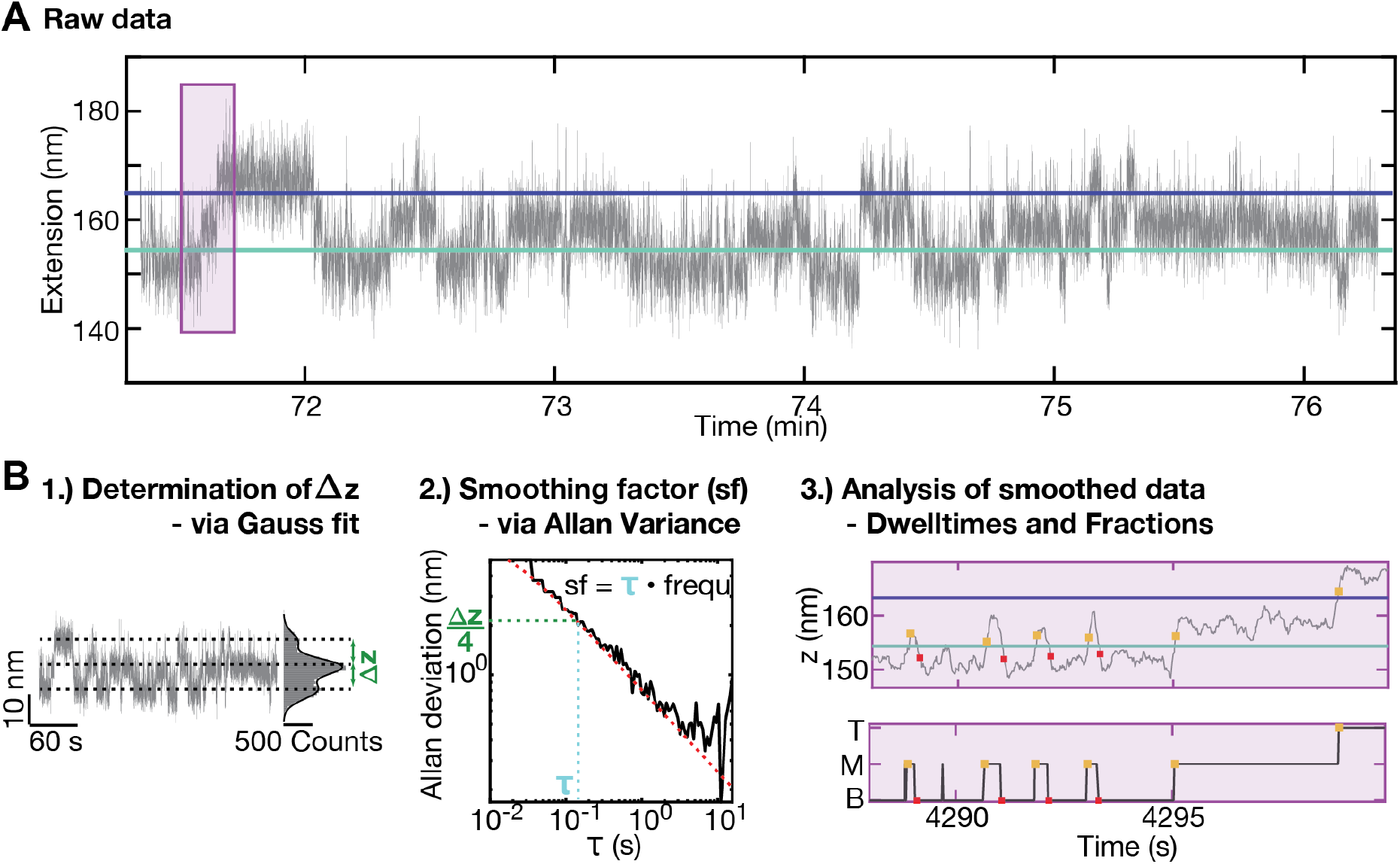
Analysis procedure of extension-time traces exhibiting three-state transitions. **A** Raw data of an extension-time trace, recorded at 58 Hz in one constant force plateau. Thresholds separating bottom (B), middle (M), and top (T) states are shown as green and blue solid lines. Noise of the trace induces transitions over the thresholds in addition to transitions caused by domain opening and closing. The purple box indicates exemplary segment of the trace analyzed in panel B3 after the analysis. **B** Procedure to determine smoothing factor for dwell time analysis. In a first step, a three-term Gaussian is fit to the extension histogram and the distance between the fitted peaks indicates the Δz that needs to be resolved for dwell time analysis. Dashed lines indicate the T, M, and B state. Secondly, the Allan deviation for a 30 second time interval at the lowest force plateau for this molecule is calculated (solid black line, according to Equation 1) and fit with a theoretical model of the Allan deviation (red dashed line). The Allan deviation is defined as the square root of one-half of the averaged square distance between the means of neighbouring intervals of length τ. Intuitively, it gives a measure of the spatial resolution after averaging over a time interval τ. Using the criterion that the deviation should be four times smaller than the distance Δz (green dashed line), a smoothing factor for each trace is determined depending on its noise level (blue dashed line). This smoothing factor is applied to the trace before analysing dwell times and fractions in a third step (right panel). Yellow squares indicate the first data point after crossing the threshold from below, i.e. transition from B to M or from M to T; red squares indicate the first data point after crossing the threshold from above, i.e. transition from T to M or M to B.

**Supplementary Figure S3.**
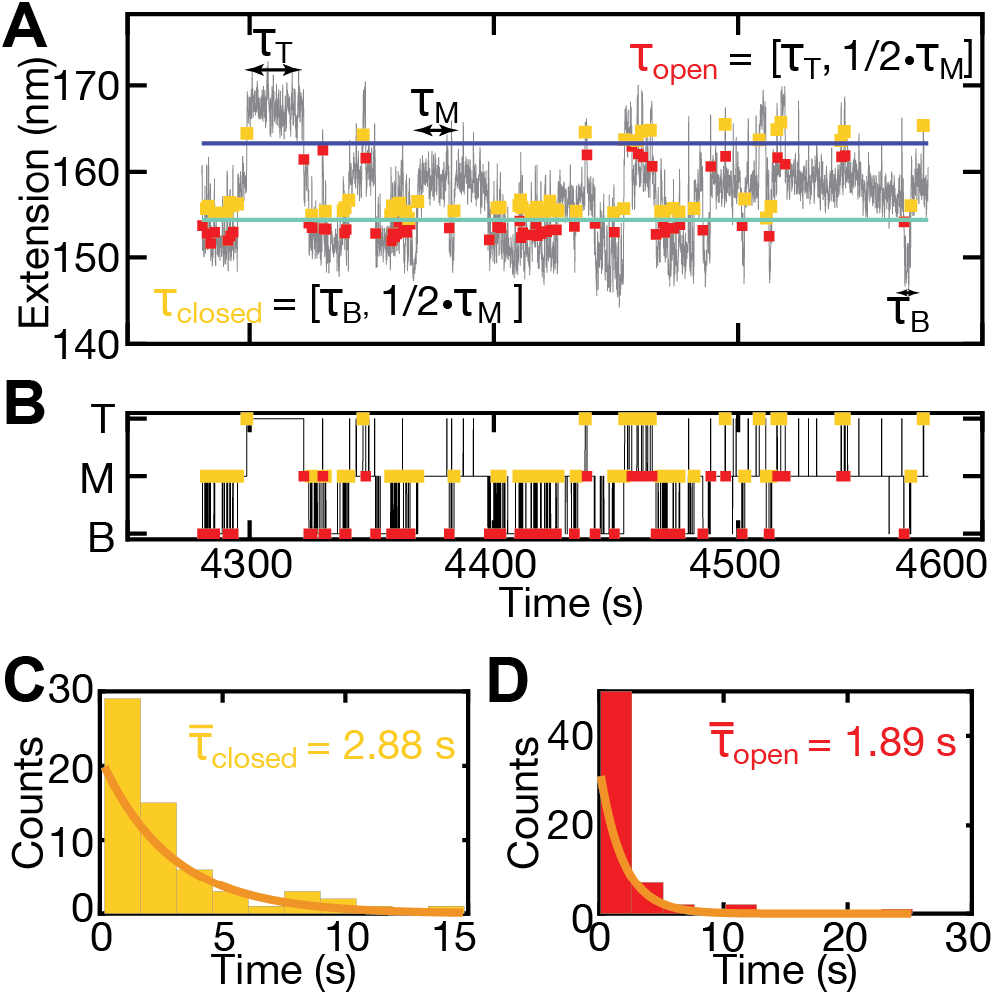
Exemplary dwell time evaluation for an extension-time trace at constant force. **A** Short segment of a extension-time trace measured for a wild type D’D3 domain VWF dimer exhibiting three-state transitions at a force of 8.4 pN. Raw data are filtered with an 11-frame moving average (which is the smoothing factor determined according to Supplementary Figure 2 for this molecule). The green horizontal line is the threshold between the bottom state (B) and the middle state (M) and the blue horizontal line is the threshold between the middle state (M) and the top state (T); yellow squares indicate the first data point after crossing the threshold from below, i.e. transition from B to M or from M to T; red squares indicate the first data point after crossing the threshold from above, i.e. transition from T to M or M to B. **B** Time trace derived from the analysis shown in panel A, indicating the current state of D3 domains with “T” corresponding to both domains opened, “M” corresponding to one domain open and one domain closed and “B” corresponding to both domains closed. The time between the transitions between “B” and “M” and “M” and “T” correspond to the dwell times. To obtain pseudo dwell time distributions of the individual domains, dwell times in the bottom state were collected together with half of the dwell times in the middle state for a distribution of τ_closed_ and dwell times in the top state were collected with half of the dwell times in the middle state for a distribution of τ_open_. **C, D** Histograms of pseudo dwell time distribution in the closed state (**C**) and the open state (**D**) obtained from the analysis shown in panels A and B. The pseudo dwell times are well described by single exponential fits, shown as solid orange line. Insets show the mean dwell time.

**Supplementary Figure S4.**
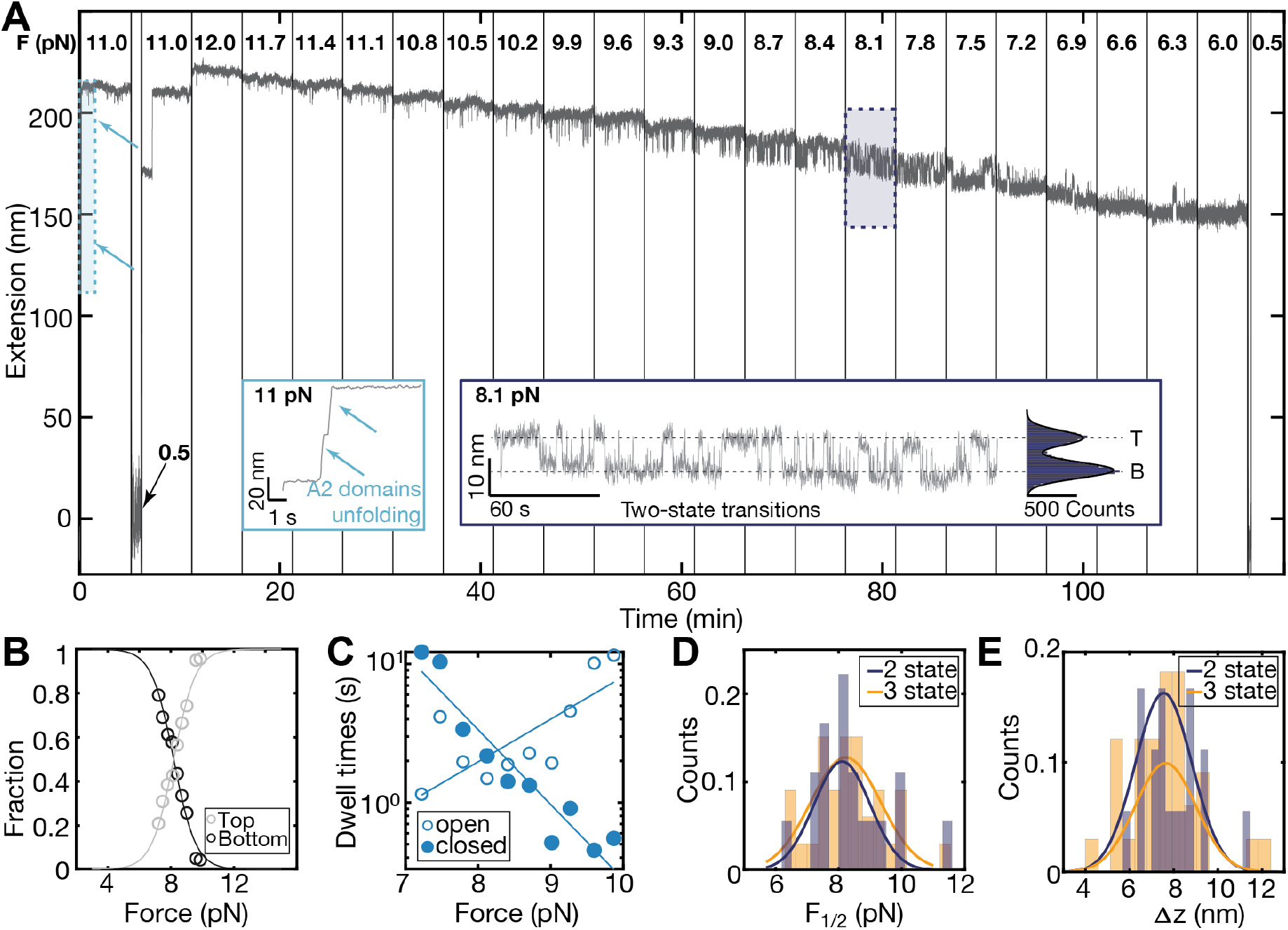
Two-state transitions in a wild type VWF dimer construct. **A** Extension-time trace for a VWF dimer showing only transitions between two states. The same experimental force ramp was conducted as for molecules exhibiting three-state transitions (see Figure 2). Insets show two the A2 unfolding events that are used as a molecular fingerprint, indicating a specifically attached dimer (blue inset), and two-state transitions close to the midpoint force at 8.1 pN together with a histogram of the extension (dark blue inset). Top and Bottom states are indicated with dashed lines. Extension was smoothed with a 9-frame moving average filter. **B** Two-state model fit to the fraction of time spent in the top and in the bottom state. Fit parameters are the midpoint force and the distance between the states. **C** Dwell time distributions in the top and in the bottom state. Unlike for the three-state transitions, dwell times for an individual domain can be determined directly for two-state transitions. **D** Distributions of midpoint forces evaluated from fitting two- and three-state transition molecules, respectively. Solid lines are Gaussian fits. Distributions are equal within experimental error. Fit parameters (95 % confidence bounds) are: Mean ± std. are: F_1/2, 3-state_ = 8.2 ± 0.9 pN and F_1/2, 2-state_ = 8.4 ± 1.2 pN. **E** Distributions of Δz values evaluated from fitting two- and three-state transition molecules respectively. Solid lines are Gaussian fits. Distributions are equal within experimental error. Mean ± std. are: Δz_3-state_ = 7.1 ± 2.0 nm and Δz_2-state_ = 8.0 ± 1.3 nm. Distributions in D and E include 15 three-state transition molecules and 18 two-state transition molecules.

**Supplementary Figure S5.**
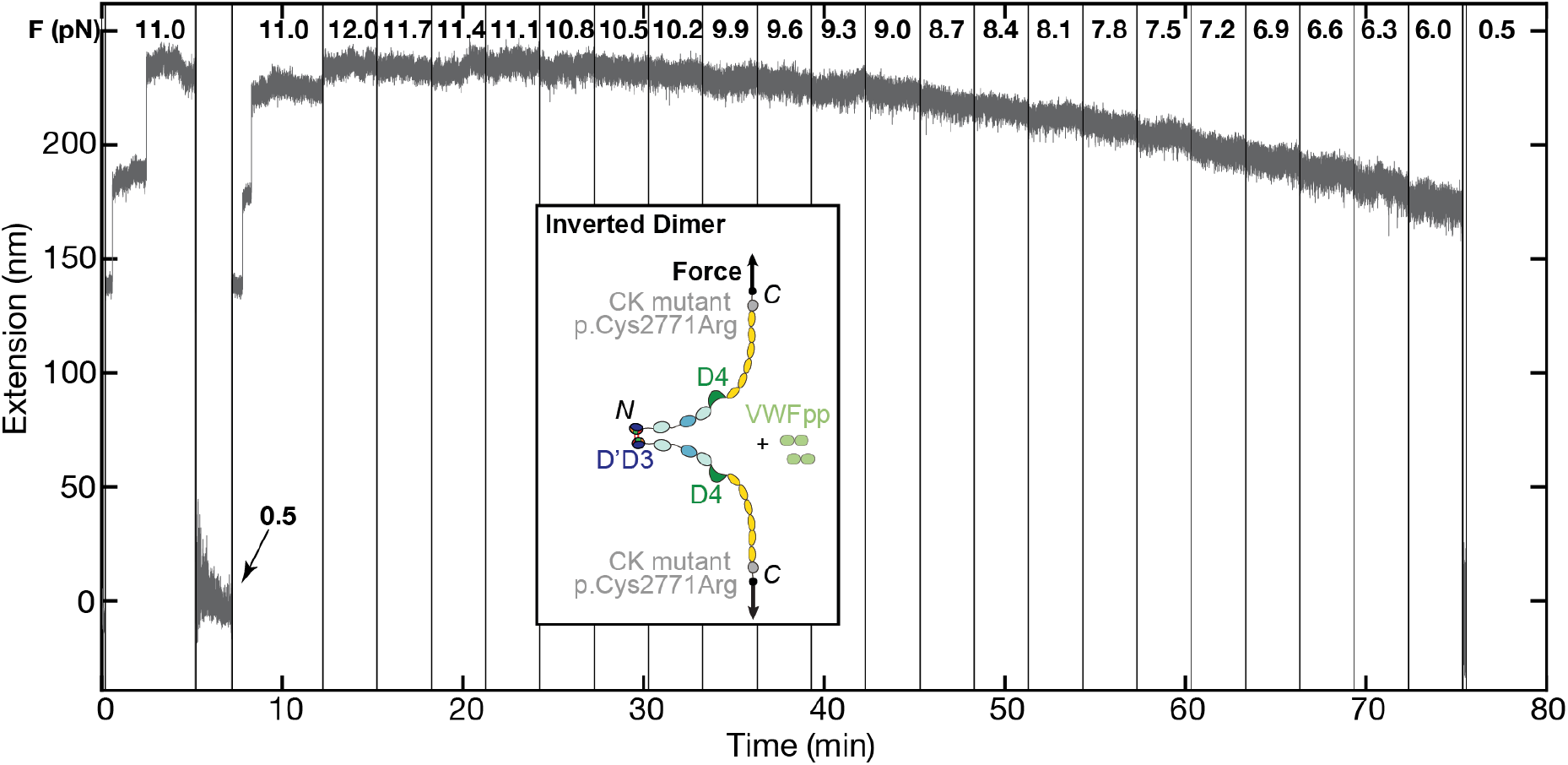
Physiological force-loading of an inverted dimer. Extension-time traces under the same force ramp protocol that was used to analyze the two- and three-state transitions with an inverted dimer (schematic of construct shown as inset). In this conformation, force propagates through the D’D3 domain as through VWF that is multimerized by cysteine linkage in the D’D3 domains (see Figure 1). Extension was smoothed with a 5-frame moving average filter. In total, >40 inverted molecules with two A2 unfolding events were screened for transitions, but none showed any.

### Supplementary Tables

**Supplementary Table S1:** Buffers used for MT measurements. The pH of buffers was adjusted with HCl and NaOH. For the measurement, buffers were supplemented with 0.1 % Tween-20 to reduce unspecific interactions of beads.

| Name | Ingredients |
| --- | --- |
| pH 7.4 | 20 mM Hepes, 150 mM NaCl, 1 mM CaCl<sub>2</sub>, 1 mM MgCl<sub>2</sub> |
| pH 7.4 + EDTA | 20 mM Hepes, 150 mM NaCl, 10 mM EDTA |
| pH 6.2 | 20 mM BisTris, 150 mM NaCl, 1 mM CaCl<sub>2</sub>, 1 mM MgCl<sub>2</sub> |
| pH 5.5 | 20 mM Na-Acetate, 150 mM NaCl, 1 mM CaCl<sub>2</sub>, 1 mM MgCl<sub>2</sub> |

## References

1. Löf, A., Müller, J. P. & Brehm, M. A. A biophysical view on von Willebrand factor activation. J. Cell. Physiol. 233, 799–810 (2018).

2. Fu, H. et al. Flow-induced elongation of von Willebrand factor precedes tension-dependent activation. Nat. Commun. 8, (2017).

3. Li, F., Li, C. Q., Moake, J. L., López, J. A. & McIntire, L. V. Shear stress-induced binding of large and unusually large von Willebrand factor to human platelet glycoprotein Ibα. Ann. Biomed. Eng. 32, 961–969 (2004).

4. Zhou, Y. F. et al. Sequence and structure relationships within von Willebrand factor. Blood 120, 449–458 (2012).

5. Springer, T. A. Von Willebrand factor, Jedi knight of the bloodstream. Blood 124, 1412–1425 (2014).

6. Zhang, X., Halvorsen, K., Zhang, C. Z., Wong, W. P. & Springer, T. A. Mechanoenzymatic cleavage of the ultralarge vascular protein von willebrand factor. Science 324, 1330–1334 (2009).

7. Huang, R. et al. Assembly of Weibel – Palade body-like tubules from N-terminal domains of von Willebrand factor. Proc. Natl. Acad. Sci. 105, 482–487 (2008).

8. Wagner, D. D. Cell Biology of von Willebrand Factor. Annu. Rev. Cell Biol. 6, 217–242 (1990).

9. Lenting, P. J., Christophe, O. D. & Denis, C. V. Von Willebrand factor biosynthesis, secretion, and clearance: Connecting the far ends. Blood 125, 2019–2028 (2015).

10. Müller, J. P. et al. pH-Dependent Interactions in Dimers Govern the Mechanics and Structure of von Willebrand Factor. Biophys. J. 111, 312–322 (2016).

11. Springer, T. A. Biology and physics of von Willebrand factor concatamers. J. Thromb. Haemost. 9, 130–143 (2011).

12. Zhou, Y. F. et al. A pH-regulated dimeric bouquet in the structure of von Willebrand factor. EMBO J. 30, 4098–4111 (2011).

13. Wagner, D. D. & Marder, V. J. Biosynthesis of von Willebrand protein by human endothelial cells: Processing steps and their intracellular localization. J. Cell Biol. 99, 2123–2130 (1984).

14. Purvis, A. R. et al. Two Cys residues essential for von Willebrand factor multimer assembly in the Golgi. Proc. Natl. Acad. Sci. U. S. A. 104, 15647–15652 (2007).

15. Dong, Z. et al. Disulfide bonds required to assemble functional von Willebrand factor multimers. J. Biol. Chem. 269, 6753–6758 (1994).

16. Dong, X. et al. The von Willebrand factor D’D3 assembly and structural principles for factor VIII binding and concatemer biogenesis. Blood 133, 1523–1533 (2019).

17. Fuller, J. R., Knockenhauer, K. E., Leksa, N. C., Peters, R. T. & Batchelor, J. D. Molecular determinants of the factor VIII/von Willebrand factor complex revealed by BIVV001 cryo-electron microscopy. Blood 137, 2970–2980 (2021).

18. Dasgupta, S. et al. VWF protects FVIII from endocytosis by dendritic cells and subsequent presentation to immune effectors. Blood 109, 610–612 (2007).

19. Lillicrap, D. von Willebrand disease: advances in pathogenetic understanding, diagnosis, and therapy. Hematology Am. Soc. Hematol. Educ. Program 2013, 254–260 (2013).

20. Shiltagh, N. et al. Solution structure of the major factor VIII binding region on von Willebrand factor. Blood 123, 4143–4151 (2014).

21. Yee, A. et al. Visualization of an N-terminal fragment of von Willebrand factor in complex with factor VIII. Blood 126, 939–942 (2015).

22. Chiu, P. L. et al. Mapping the interaction between factor VIII and von Willebrand factor by electron microscopy and mass spectrometry. Blood 126, 935–938 (2015).

23. Strick, T. R., Allemand, J. F., Bensimon, D., Bensimon, A. & Croquette, V. The elasticity of a single supercoiled DNA molecule. Science (80-.). 271, 1835–1837 (1996).

24. Vilfan, I. D., Lipfert, J., Koster, D. A., Lemay, S. G. & Dekker, N. H. Magnetic Tweezers for Single-Molecule Experiments. in Handbook of Single-Molecule Biophysics (eds. Hinterdorfer, P. & Oijen, A.) 371–395 (Springer US, 2009). doi:10.1007/978-0-387-76497-9-13

25. Kriegel, F., Vanderlinden, W., Nicolaus, T., Kardinal, A. & Lipfert, J. Measuring single-molecule twist and torque in multiplexed magnetic tweezers. Methods Mol. Biol. 1814, 75–98 (2018).

26. Löf, A. et al. Multiplexed protein force spectroscopy reveals equilibrium protein folding dynamics and the low-force response of von Willebrand factor. Proc. Natl. Acad. Sci. U. S. A. 116, 18798–18807 (2019).

27. Le, S., Liu, R., Lim, C. T. & Yan, J. Uncovering mechanosensing mechanisms at the single protein level using magnetic tweezers. Methods 94, 13–18 (2016).

28. Popa, I. et al. A HaloTag Anchored Ruler for Week-Long Studies of Protein Dynamics. J. Am. Chem. Soc. 138, 10546–10553 (2016).

29. Ott, W. et al. Elastin-like Polypeptide Linkers for Single-Molecule Force Spectroscopy. ACS Nano 11, 6346–6354 (2017).

30. Gruber, S. et al. Designed anchoring geometries determine lifetimes of biotin-streptavidin bonds under constant load and enable ultra-stable coupling. Nanoscale 12, 21131–21137 (2020).

31. Müller, J. P. et al. Force sensing by the vascular protein von Willebrand factor is tuned by a strong intermonomer interaction. Proc. Natl. Acad. Sci. U. S. A. 113, 1208–1213 (2016).

32. Ainavarapu, S. R. K. et al. Contour length and refolding rate of a small protein controlled by engineered disulfide bonds. Biophys. J. 92, 225–233 (2007).

33. Cao, Y., Kuske, R. & Li, H. Direct observation of Markovian behavior of the mechanical unfolding of individual proteins. Biophys. J. 95, 782–788 (2008).

34. Chen, H. et al. Dynamics of Equilibrium Folding and Unfolding Transitions of Titin Immunoglobulin Domain under Constant Forces. J. Am. Chem. Soc. 137, 3540–3546 (2015).

35. Bell, G. I. Models for the specific adhesion of cells to cells. Science (80-.). 200, 618–627 (1978).

36. Dong, X. & Springer, T. A. Disulfide Exchange in von Willebrand factor Dimerization in the Golgi. Blood 137, 1263–1267 (2020).

37. Phan, J. et al. Structural basis for the substrate specificity of tobacco etch virus protease. J. Biol. Chem. 277, 50564–50572 (2002).

38. Theile, C. S. et al. Site-specific N-terminal labeling of proteins using sortase-mediated reactions. Nat. Protoc. 8, 1800–1807 (2013).

39. Brehm, M. A. et al. von Willebrand disease type 2A phenotypes IIC, IID and IIE: A day in the life of shear-stressed mutant von Willebrand factor. Thromb. Haemost. 112, 96–108 (2014).

40. Lippok, S. et al. Von Willebrand factor is dimerized by protein disulfide isomerase. Blood 127, 1183–1191 (2016).

41. Dong, X. et al. The von Willebrand factor D’D3 assembly and structural principles for factor VIII binding and concatemer biogenesis. Blood 133, 1523–1533 (2019).

42. Humphrey, W., Dalke, A. & Schulten, K. VMD: Visual molecular dynamics. J. Mol. Graph. 14, 33–38 (1996).

43. Ribeiro, J. V. et al. QwikMD - Integrative Molecular Dynamics Toolkit for Novices and Experts. Sci. Rep. 6, 1–14 (2016).

44. Best, R. B. et al. Optimization of the additive CHARMM all-atom protein force field targeting improved sampling of the backbone φ, ψ and side-chain χ1 and χ2 Dihedral Angles. J. Chem. Theory Comput. 8, 3257–3273 (2012).

45. MacKerell, A. D. et al. All-atom empirical potential for molecular modeling and dynamics studies of proteins. J. Phys. Chem. B 102, 3586–3616 (1998).

46. Walker, P. U., Vanderlinden, W. & Lipfert, J. Dynamics and energy landscape of DNA plectoneme nucleation. Phys. Rev. E 98, 42412 (2018).

47. Lipfert, J., Hao, X. & Dekker, N. H. Quantitative modeling and optimization of magnetic tweezers. Biophys. J. 96, 5040–5049 (2009).

48. Cnossen, J. P., Dulin, D. & Dekker, N. H. An optimized software framework for realtime, high-throughput tracking of spherical beads. Rev. Sci. Instrum. 85, 103712 (2014).

49. Te Velthuis, A. J. W., Kerssemakers, J. W. J., Lipfert, J. & Dekker, N. H. Quantitative guidelines for force calibration through spectral analysis of magnetic tweezers data. Biophys. J. 99, 1292–1302 (2010).

50. de Vlaminck, I., Henighan, T., van Loenhout, M. T. J., Burnham, D. R. & Dekker, C. Magnetic forces and dna mechanics in multiplexed magnetic tweezers. PLoS One 7, (2012).

51. Ostrofet, E., Papini, F. S. & Dulin, D. Correction-free force calibration for magnetic tweezers experiments. Sci. Rep. 8, 1–10 (2018).

52. Ott, W. et al. Elastin-like Polypeptide Linkers for Single-Molecule Force Spectroscopy. ACS Nano 11, 6346–6354 (2017).

53. Yin, J. et al. Genetically encoded short peptide tag for versatile protein labeling by Sfp phosphopantetheinyl transferase. Proc. Natl. Acad. Sci. U. S. A. 102, 15815–15820 (2005).

54. Chen, I., Dorr, B. M. & Liu, D. R. A general strategy for the evolution of bond-forming enzymes using yeast display. Proc. Natl. Acad. Sci. U. S. A. 108, 11399–11404 (2011).

55. Durner, E., Ott, W., Nash, M. A. & Gaub, H. E. Post-Translational Sortase-Mediated Attachment of High-Strength Force Spectroscopy Handles. ACS Omega 2, 3064–3069 (2017).

56. Allan, D. W. Statistics of Atomic Frequency Standards. Proc. IEEE 54, 221–230 (1966).

57. Czerwinski, F., Richardson, A. C. & Oddershede, L. B. Quantifying Noise in Optical Tweezers by Allan Variance. Opt. Express 17, 13255 (2009).

58. van Oene, M. M. et al. Quantifying the Precision of Single-Molecule Torque and Twist Measurements Using Allan Variance. Biophys. J. 114, 1970–1979 (2018).

